# Role of anterior insula cortex in context-induced relapse of nicotine-seeking

**DOI:** 10.1101/2021.12.08.471717

**Authors:** Hussein Ghareh, Isis Alonso-Lozares, Dustin Schetters, Rae J. Herman, Tim S. Heistek, Yvar van Mourik, Philip Jean-Richard-dit-Bressel, Gerald Zernig, Huibert D. Mansvelder, Taco J. de Vries, Nathan J. Marchant

**Author notes:** Corresponding Author: Nathan Marchant. denotes co-first author (Hussein Ghareh, Isis Alonso-Lozares).

## Abstract

Tobacco use is the leading cause of preventable death worldwide, and relapse during abstinence remains the key barrier to successful treatment of tobacco addiction. During abstinence, environmental contexts associated with nicotine use can induce craving and contribute to relapse. The insular cortex (IC) is thought to be a critical substrate of nicotine addiction and relapse. However, its specific role in context-induced relapse of nicotine-seeking is not fully known. In this study, we report a novel rodent model of context-induced relapse to nicotine-seeking after punishment-imposed abstinence, which models self-imposed abstinence through increasing negative consequences of excessive drug use. Using the neuronal activity marker Fos we find that the anterior (aIC), but not the middle or posterior IC, shows increased activity during context-induced relapse. Combining Fos with retrograde labelling of aIC inputs, we show projections to aIC from contralateral aIC and basolateral amygdala exhibit increased activity during context-induced relapse. Next, we used fiber photometry in aIC and observed phasic increases in aIC activity around nicotine-seeking responses during self-administration, punishment, and the context-induced relapse tests. Next, we used chemogenetic inhibition in both male and female rats to determine whether activity in aIC is necessary for context-induced relapse. We found that chemogenetic inhibition of aIC decreased context-induced nicotine-seeking after either punishment- or extinction-imposed abstinence. These findings highlight the critical role nicotine-associated contexts play in promoting relapse, and they show that aIC activity is critical for this context-induced relapse following both punishment and extinction imposed abstinence.

## Introduction

Tobacco use is one of the leading causes of preventable death worldwide. In both abstinent and non-abstinent individuals with a history of nicotine use, exposure to cues associated with nicotine use provokes craving (1, 2), which is strongly related to relapse (3). Environmental contexts also play a crucial role in nicotine craving. An environmental context associated with nicotine use retains the ability to reinstate cue-induced nicotine craving after extinction in humans (4, 5). Pre-clinical models have been used to study the role of contexts in relapse using the extinction-based context-induced reinstatement (or ABA renewal) model (6, 7). One potential limitation of the extinction-based models is that extinction does not capture the motivation for abstinence in humans (8, 9). We recently developed a variation of this model in which an alcohol-reinforced response is suppressed by response-contingent punishment (10). These studies built on prior models using punishment to model the negative consequences of drug use (11–14). We and others have demonstrated context-induced relapse of alcohol, food, and cocaine seeking after punishment-imposed abstinence in an alternative context (15–21). The extent to which this phenomenon translates to context-induced relapse of nicotine-seeking has not yet been demonstrated.

The insular cortex (IC) has been considered a critical neural substrate of nicotine addiction since it was discovered that some human patients with stroke-induced damage to their insula had a higher probability of smoking cessation (22). Subsequent clinical studies found that nicotine dependence is positively correlated with cue-induced activation in the insula (23, 24), and there is a negative association between nicotine dependence and insula structural integrity (25). Insula activity is related to the processing of drug cues (26), and cue-induced activity in anterior insula is indicative of relapse vulnerability (27). Both nicotine withdrawal and acute abstinence lead to changes in anterior insula activity, and can also weaken connectivity between the default mode network and salience network at the resting state (28, 29). In light of these findings, we focus here on the role of the rodent anterior insula cortex (aIC) in context-induced relapse of punished nicotine-seeking.

Here we demonstrate for the first time, in both male and female rats, context-induced relapse of nicotine-seeking after punishment of nicotine taking in an alternative context. Using the neuronal marker of activity Fos (30–32), we show that context-induced relapse of punished nicotine-seeking is associated with increased Fos expression in aIC but not middle or posterior IC. We also found that context-induced relapse was associated with increased Fos in projections to aIC from both contralateral aIC and ipsilateral basolateral amygdala (BLA). Next, we used calcium imaging with fiber photometry (33, 34) to record the activity of aIC neurons throughout nicotine self-administration, punishment, and context-induced relapse. We found that aIC activity was associated with both nicotine infusion and punishment, and also nicotine seeking during the relapse test. To determine a causal role for activity in aIC and context induced relapse, we used chemogenetics (35) to inhibit activity in aIC, and found that this decreased context-induced relapse after punishment. Because of potential differences in the neural control of context-induced relapse after punishment or extinction (36), we next tested chemogenetic inhibition of aIC after extinction. We also found that this inhibition decreased context-induced reinstatement of nicotine seeking. These data highlight the critical role that nicotine-associated contexts play in promoting relapse, and they show that activity in aIC is necessary for this, further highlighting a critical role of this structure in relapse to nicotine use.

## Results

### Exp. 1: Context-induced relapse to nicotine-seeking after punishment-imposed abstinence is associated with increased Fos expression in aIC, and projections from BLA to aIC

#### Behavioral data

Statistical analysis of the training data (Fig. 1B) revealed a significant Nose-Poke x Session interaction (F(14,210) = 16.1, p < 0.001), indicating that responses on the active nose-poke increased throughout training compared to inactive nose-pokes. In punishment (Fig. 1C) we observed a significant Nose-poke x Session interaction (F(6,90) = 9.6; p < 0.001), reflecting the decrease in active nose-pokes during punishment. On the final test we returned the rats to either context B (Punishment) or context A (Nicotine) (Fig. 1D). We found a significant main effect of Test Context (F(1,13) = 14.7; p < 0.01), and a Test Context x Nose-poke interaction (F(1,13) = 6.4; p < 0.05). These data show that rats tested in the nicotine context significantly increased nicotine-seeking (active nose-pokes) compared to the rats tested in the punishment context.

**Figure 1.**
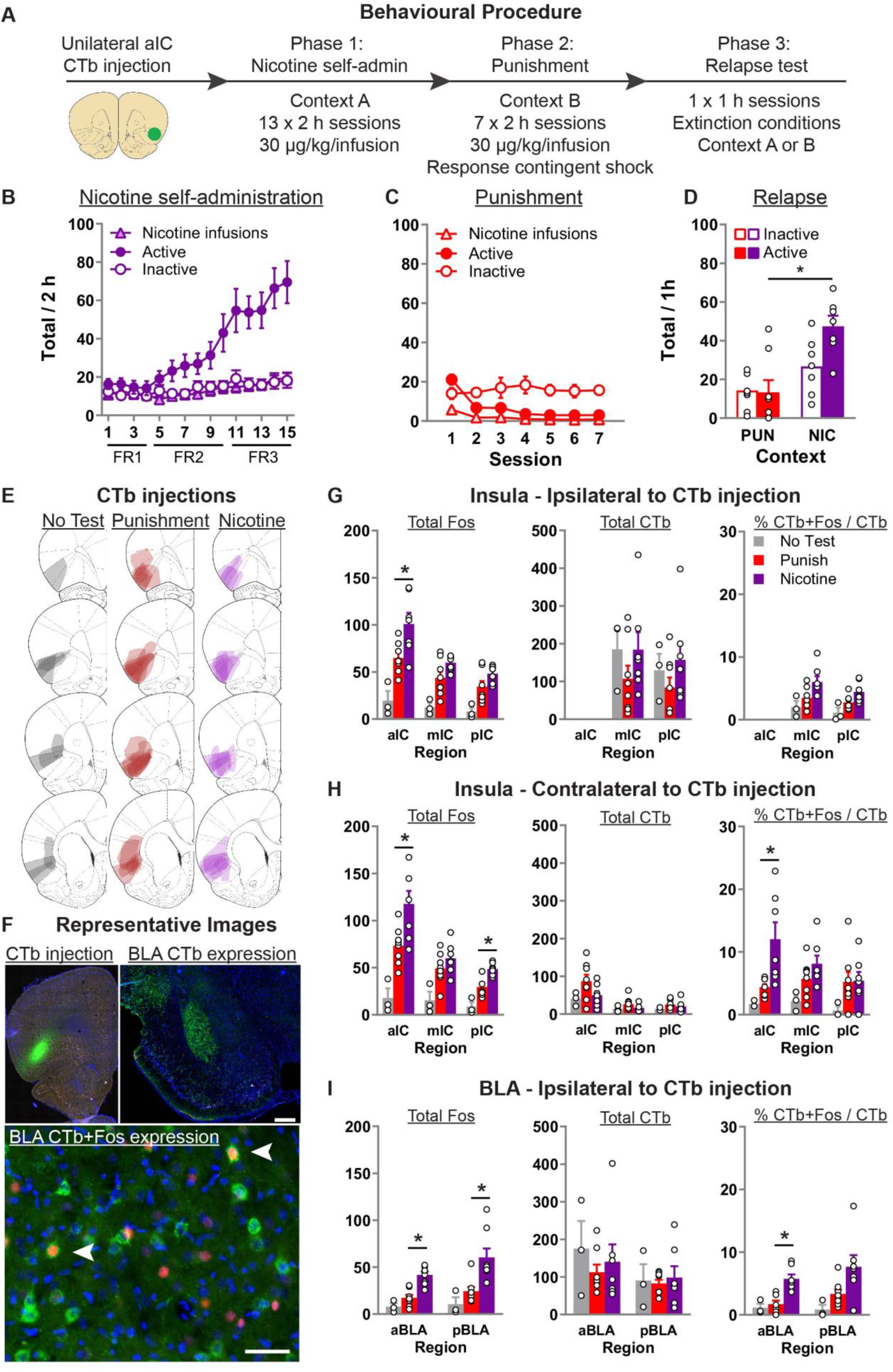
*Context-induced relapse of punished nicotine-seeking is associated with selective activation of BLA→aIC and contralateral aIC→aIC projections.* (**A**) Outline of the experimental procedure (n = 19 female). (**B, C, D**) Mean±sem active and inactive nose-pokes, and nicotine infusions, during nicotine self-administration in context A (**B**), punishment in context B (**C**), and the context-induced nicotine-relapse test in context B or A (**D**). (**E**) Representative plots of the spread of CTb injections for the three groups. (**F**) Representative images of CTb injection in aIC, and CTb+Fos in BLA. (**G, H, I**) Data are mean ± SEM number of Fos or CTb neurons per mm^2^, and percentage Ctb + Fos neurons, in the IC hemisphere ipsilateral to CTb injection (**G**), IC hemisphere contralateral to the CTb injection (**H**), or BLA ipsilateral to the CTb injection (**I**). *p < 0.05; aIC, anterior insula cortex; mIC, middle insula cortex; pIC, posterior insula cortex; BLA, Basolateral Amygdala; FR, fixed-ratio.

#### CTb+Fos data

Fig. 1E shows the spread of CTb at injection site, Fig. 1F shows example CTb injection in aIC, CTb labelling in pIC and BLA, and CTb+Fos neurons in BLA. Fig. 1G shows the total Fos, CTb, and percentage CTb+Fos neurons in the insula cortex *ipsilateral* to CTb injection. Using separate one-way ANOVA for each region, we found a significant effect of Test Context for Total Fos in aIC (F(2,17) = 13.1; p = 0.001), mIC (F(2,17) = 12.6; p = 0.001), and pIC (F(2,17) = 10.1; p = 0.002). Subsequent Tukey post-hoc only revealed a significant difference between rats tested in nicotine versus punishment context for aIC (p = 0.025). We found no effect of Test Context for the total CTb in mIC and pIC (Fs < 1; ps > 0.05). For percent of CTb neurons that also express Fos, there was a main effect of Test Context in both mIC (F(2,17) = 5.3; p < 0.05) and pIC (F(2,17) = 6.5; p < 0.05). However post-hoc analysis revealed no significant difference between the Nicotine and Punishment tested rats.

Fig. 1H shows the total Fos, CTb, and percentage CTb+Fos neurons in the insula cortex *contralateral* to CTb injection. Using separate one-way ANOVA for each region, we found a comparable pattern of effects for Total Fos. Specifically, there was a main effect of Test Context in aIC (F(2,17) = 14.1; p < 0.001), mIC (F(2,17) = 8.3; p < 0.01), and pIC (F(2,17) = 21.6; p = 0.002), and subsequent post-hoc revealed a significant difference between rats tested in nicotine or punishment context in aIC (p = 0.02) as well as pIC (p = 0.003), but not mIC (p = 0.32). In all three regions we found no effect of Test Context on total CTb (Fs < 2.5; ps > 0.05). For the percent of CTb neurons that also express Fos, there was a main effect of Test Context in both aIC (F(2,17) = 7.4; p < 0.01) and mIC (F(2,17) = 4.2; p < 0.05), but not pIC (F(2,17) = 1.8; p > 0.05). Post-hoc analysis revealed significant difference between the Nicotine and Punishment tested rats in aIC (p = 0.01) but not mIC (p = 0.3). These data show that context-induced relapse of nicotine-seeking is associated with increased activity in the aIC neurons that project to the contralateral aIC.

In Fig. 1I we show the total Fos, total CTb, and the percentage of CTb neurons that are also Fos positive in the Basolateral Amygdala (BLA) of the *ipsilateral* hemisphere to the CTb injection. One-way ANOVA revealed a significant effect of Test Context in both anterior BLA (F(2,17) = 21.1; p < 0.001) and posterior BLA (F(2,17) = 9.4; p < 0.001). Subsequent Tukey post-hoc revealed a significant difference between rats tested in nicotine or punishment context in aBLA (p < 0.001) and pBLA (p = 0.007). We found no effect of Test Context on total CTb in both aBLA (F(2,17) < 1; p > 0.05) and pBLA (F(2,17) < 1; p > 0.05). Finally, analysis of the percent of CTb positive neurons that are also Fos positive revealed a significant effect of Test Context in aBLA (F(2,17) = 13.5; p < 0.001) and pBLA (F(2,17) = 4.6; p < 0.05). Tukey post-hoc analysis revealed a significant difference between rats tested in nicotine versus punishment context in aBLA (p = 0.001) but not pBLA (p = 0.1). In summary, these data show that context-induce relapse of nicotine-seeking is associated with increased activity in BLA, and that there is also selectively increased activity in the aBLA→aIC pathway.

### Exp. 2: Real-time neuronal activity in aIC encodes nicotine-seeking responses across nicotine self-administration, punishment, and context-induced relapse

We examined real-time population-level aIC principal neuron calcium (Ca2+) transients throughout the entire task (Fig. 2A). We used AAV encoding jGCaMP7f under control of the hSyn1 promoter to express the Ca2+ sensor jGCaMP7f (37) in aIC. Fluorescence was measured via an optic fiber cannula implanted in aIC (Fig. 2B). We analyzed Ca2+ transients around nosepokes using 95% confidence intervals with a consecutive threshold of 0.25 s (38).

**Figure 2.**
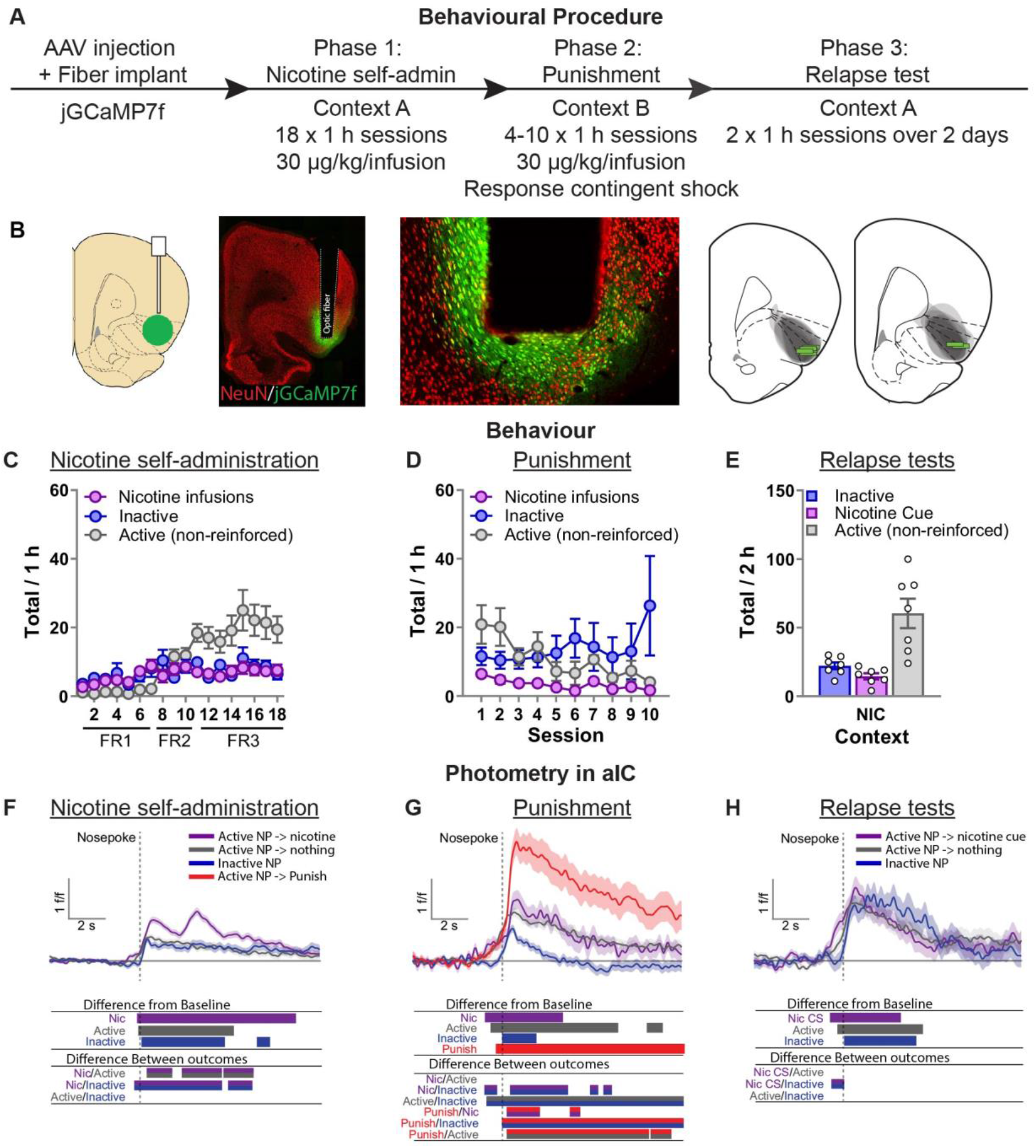
*Photometry reveals nicotine and punishment-associated activity in aIC.* (**A**) Outline of the experimental procedure (n = 7 female). (**B**) Representative images of jGCaMP7f expression and fiber implant in aIC. (**C, D, E**) Mean±sem nicotine infusions, inactive nose-pokes, and non-reinforced active nose-pokes during nicotine self-administration in context A (**C**), punishment in context B (**D**), and the context-induced nicotine-seeking tests (**E**). (**F**) Ca2+ traces around the nose-poke in aIC self-administration in context A (Reinforced active (Nic): n = 751; Non-reinforced active: n = 838; Inactive: n = 588). (**G**) Ca2+ traces around the nose-poke in aIC during punishment in context B (Reinforced active (Nic): n = 89; Non-reinforced active: n = 366; Inactive: n = 318; Punish: n = 137). (**H**) Ca2+ traces around the nose-poke in aIC during context-induced relapse test in context A (Active + Nic CS: n = 95; Active non-reinforced: n = 187; Inactive: n = 80). For all photometry traces, bars at bottom of graph indicate significant deviations from baseline (dF/F ≠ 0), or significant differences between the specific events (Nicotine infusion, non-reinforced active nose-poke, inactive nose-poke, Punishment, or Nicotine CS), determined via bootstrapped confidence intervals (95% CI), and permutation tests with alpha 0.008 and 0.01 for comparisons between punishment sessions, and self-administration and tests respectively. Vertical dashed line indicates nose-poke, horizontal line indicates baseline (dF/F = 0).

#### Behavioral data

Repeated measures ANOVA on the self-administration data (Fig. 2C) revealed a significant effect of Nose-Poke (F(1,5) = 12.8, p < 0.05). In punishment (Fig. 2D) repeated measures ANOVA revealed no effect of Nose-Poke (F(1,5) = 1.0, p > 0.05). On the final test we returned the rats to context A (Nicotine) over two test sessions in consecutive days (Fig. 2E). We found an overall effect of Nose-Poke (F(1,5) = 15.8; p < 0.01) reflecting greater responses on the active nose-poke compared to the inactive nose-poke in these test sessions.

#### Nicotine self-administration photometry data (Fig. 2F)

In self-administration, we found that aIC shows significant increased excitatory Ca2+ transients after nose-pokes, suggesting a general role of aIC for encoding response-outcome contingencies. Response-generated nicotine infusions (paired with a CS), exhibited a sustained robust biphasic excitatory Ca2+ transient. Importantly we found a significant difference between the reinforced and non-reinforced active nose-poke events, indicating increased activity in aIC specifically related to the nicotine-associated cue and nicotine infusion. Surprisingly, we also found that there was increased excitatory Ca2+ transients, relative to baseline, prior to the nose-poke for the two types of active nose-pokes (reinforced, or non-reinforced), but not inactive nose-poke. Direct comparisons between the events, using permutation tests, also revealed a significant difference between the reinforced active nose-poke and inactive nose-poke.

#### Punishment photometry data (Fig. 2G)

In punishment we again found that aIC shows excitatory transients after all nose-pokes. We found that active nose-pokes that resulted in punishment caused a significantly greater sustained excitatory Ca2+ transient compared to the other outcomes (nicotine infusion, nothing). Moreover, the pattern of activity to the non-punished nicotine infusion changed compared to self-administration, because there was no longer a significant difference between reinforced and non-reinforced active nose-pokes. Another interesting observation is that again, like self-administration, we found a significant increase in aIC activity in the lead up to active but not inactive nose-pokes. There was a significant increase relative to baseline for the three types of active nose-pokes (Punished, Nicotine infusions, non-reinforced), but there was no such increase prior to the inactive nose-pokes. Direct comparison between the events using permutations tests revealed a significant difference between the inactive nose-poke and both nicotine infusion and non-reinforced active nose-pokes, but not the punished nose-poke.

#### Context-induced relapse photometry data (Fig. 2H)

In the final two sessions we tested the rats in the original training context (context A) under extinction conditions over two consecutive days. We found significant excitatory Ca2+ transients in aIC, relative to baseline, after all nose-pokes. Like the prior phases, there was also a significant increase in activity prior to active but not inactive nose-pokes. There were no significant differences between active nose-pokes that elicited the nicotine-associated cue versus those that did not, suggesting differential activity observed in previous phases encoded the reinforcing and punishing outcomes (nicotine, shock), which were absent in the final test.

**52+**

### Exp. 4: Chemogenetic inhibition of aIC decreases context-induced relapse of punished nicotine-seeking

Fig. 4B shows nicotine self-administration in context A. We observed a significant Nose-Poke x Session interaction (F(12,288) = 13.4, p < 0.001), with responses on the active nose-poke increasing throughout training compared to inactive nose-pokes. Fig. 4C shows nicotine self-administration during punishment in context B. We observed a significant Nose-Poke x Session interaction (F(5,120) = 7.1, p < 0.001), with responses on the active nose-poke decreasing throughout punishment compared to inactive nose-pokes. Fig. 4D shows nicotine-seeking during the relapse tests. We observed a significant Group x Nose-poke interaction (F(1,24) = 13.2; p = 0.001), and a Context x Group x Nose-poke interaction (F(1,24) = 11.6; p = 0.001). We observed no overall effect of Sex (F(1,24) = 4.0; p > 0.05). Moreover, we found no Sex x Group x Nose-poke interaction (F(1,24) = 2.5; p > 0.05), nor Sex x Context x Group x Nose-poke interaction (F(1,24) = 3.6; p > 0.05). These results show that chemogenetic inhibition of aIC significantly decreases context-induced relapse of nicotine-seeking after punishment, in both male and female rats.

**Figure 3.**
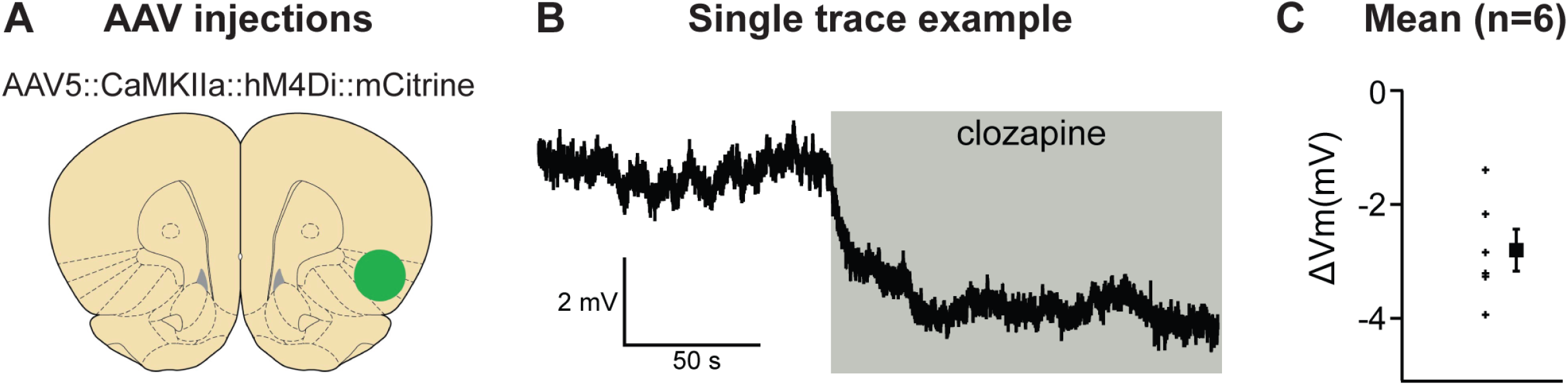
*Validation of chemogenetic inhibition of aIC neurons expressing hM4Di by clozapine.* (**A**) shows the virus injection location. (**B**) shows example trace of inward current demonstrating clear clozapine-induced hyperpolarization of the hM4Di expressing neurons. (**C**) shows the average hyperpolarization induced by clozapine for the n=6 neurons recorded (left) and the mean hyperpolarization (right) for these neurons.

**Figure 4.**
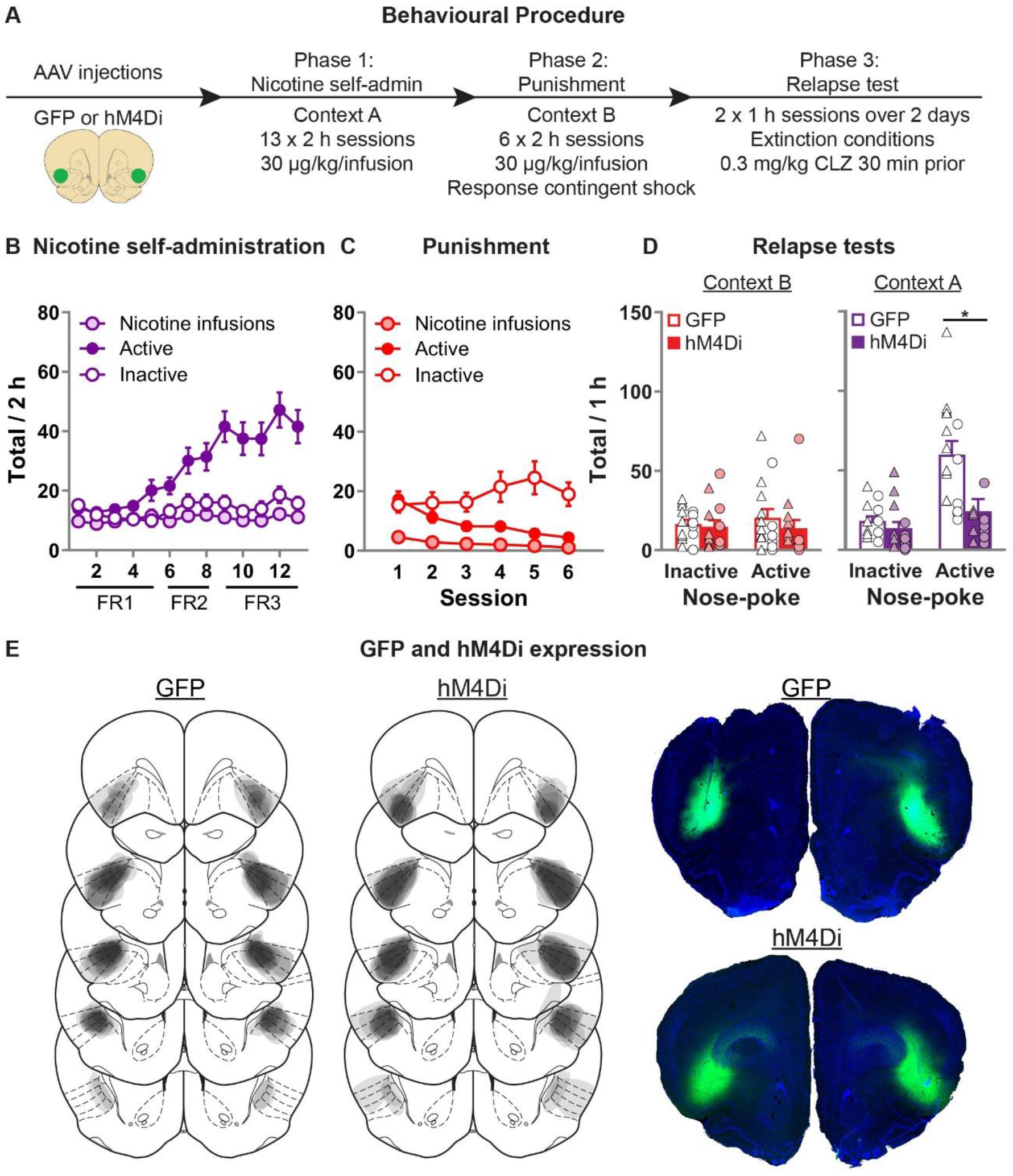
*Effect of chemogenetic inhibition of aIC on context-induced relapse of punished nicotine-seeking.* (**A**) Outline of the experimental procedure (n = 15 female, 13 male). (**B**) Mean±sem nicotine infusions, active, inactive nose-pokes during nicotine self-administration in context A. (**C**) Mean±sem nicotine infusions, active, inactive nose-pokes during punishment in context B. (**D**) Mean±sem nose-pokes during the context-induced relapse tests. Individual data also plotted, triangles = female, circles = male. (**E**) Representative plots of the spread of GFP (left) and hM4Di (middle) in aIC of rats in experiment 4. Right top shows an example section of a rat showing GFP expression in aIC, and right bottom shows an example of hM4Di expression. CLZ, Clozapine; FR, fixed-ratio.

### Exp. 5: Chemogenetic inhibition of aIC decreases context-induced relapse of extinguished nicotine-seeking

Fig. 5B shows nicotine self-administration in context A. We observed a significant Nose-Poke x Session interaction (F(12,324) = 12.3, p < 0.001), indicating that responses on the active nose-poke increased throughout training compared to inactive nose-pokes. Fig. 5C shows nicotine-seeking during extinction in context B. We observed a significant effect of Session (F(7,189) = 14.0; p < 0.001), and Nose-Poke (F(1,27) = 15.6; p < 0.001), but no Session x Nose-poke interaction (F(7,189) = 1.5; p > 0.05); both the active and inactive nose-poke decreased throughout extinction. Fig. 5D shows nicotine-seeking during the relapse tests. We observed a significant Group x Nose-poke interaction (F(1,27) = 8.5; p < 0.01), as well as a Context x Group x Nose-poke interaction (F(1,27) = 11.3; p < 0.01). We observed no overall effect of Sex (F(1,27) = 1.6; p > 0.05), and no Sex x Group x Nose-poke interaction (F(1,27) = 1.1; p > 0.05), nor Sex x Context x Group x Nose-poke interaction (F(1,24) < 1; p > 0.05). These results show that chemogenetic inhibition of aIC significantly decreases context-induced relapse of nicotine-seeking after extinction, in both male and female rats.

**Figure 5.**
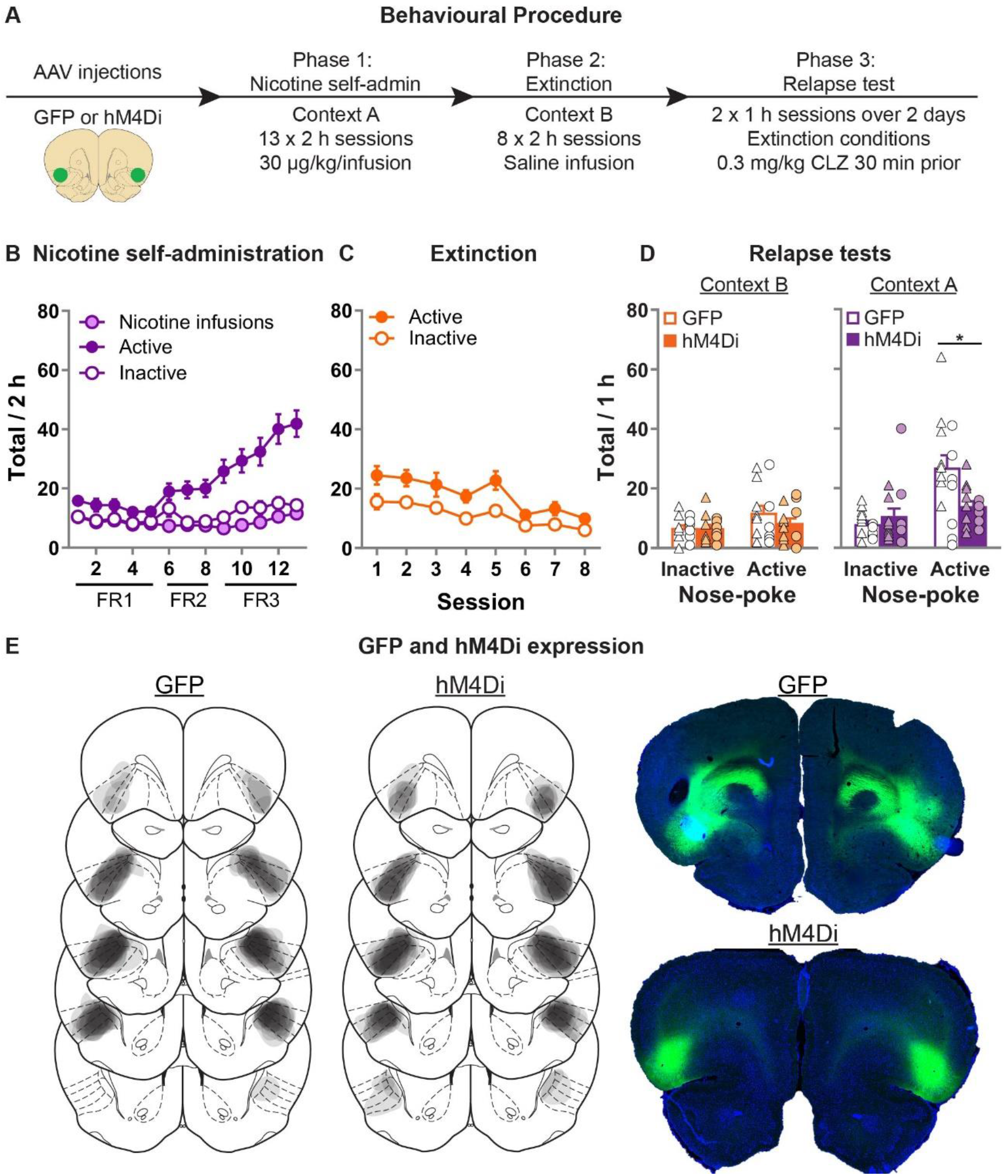
*Effect of chemogenetic inhibition of* aIC *on context-induced relapse of extinguished nicotine-seeking.* (**A**) Outline of the experimental procedure (n = 18 female, 13 male). (**B**) Mean±sem nicotine infusions, active, inactive nose-pokes during nicotine self-administration in context A. (**C**) Mean±sem active, inactive nose-pokes during extinction in context B. (**D**) Mean±sem nose-pokes during the context-induced relapse tests. Individual data also plotted, triangles = female, circles = male. (**E**) Representative plots of the spread of GFP (left) and hM4Di (middle) in aIC of rats in experiment 5. Right top shows an example section of a rat showing GFP expression in aIC, and right bottom shows an example of hM4Di expression. CLZ, Clozapine; FR, fixed-ratio.

## Discussion

In this study, we describe a novel rodent model of context-induced relapse to nicotine-seeking after punishment-imposed abstinence. We found that this form of relapse is associated with increased Fos expression in aIC, but not mIC or pIC, as well as BLA. Using retrograde tracing from aIC, we also show that inputs from contralateral aIC and ipsilateral anterior BLA are also activated during context-induced relapse of nicotine-seeking. Using fiber photometry, we found that nicotine infusions during self-administration elicited phasic increases in aIC activity. During punishment, phasic increases in aIC activity was significantly greater for the punishment outcome compared to nicotine infusion and non-reinforced responses. During the final tests we found increased activity associated with each response type. Interestingly, we also found that aIC activity increased prior to active but not inactive nose-pokes across each phase of the experiment, indicating a potential role of aIC in anticipating outcomes prior to actions, and/or selectively promoting reinforced responses. Next, we used chemogenetics to show that inhibition of aIC decreased context-induced relapse of nicotine-seeking after both punishment-imposed abstinence and after extinction. The chemogenetic experiments we conducted in both male and female rats, and we observed a comparable effect of chemogenetic inhibition of aIC in both sexes. This study demonstrates the importance of nicotine-associated contexts in promoting relapse. We show that the aIC activity is critical for this effect, regardless of the mode of abstinence, further highlighting the important role of aIC in relapse of nicotine use.

### Methodological considerations

Several issues must be considered in the interpretation of these findings. In this study, we used intravenous nicotine during self-administration. While human nicotine use is primarily through cigarette smoking, intravenous administration of nicotine is reinforcing in humans (39), demonstrating that a comparable route of administration supports reinforcement in humans. In addition, smokeless tobacco is also addictive and is responsible for many adverse health consequences (40). Recently it has been shown that vapor administration of nicotine is effective in rodents (41, 42), and vapor exposure causes both physical (43) and psychological effects (41, 44). The extent to which the route of self-administration changes the neurobiological substrates of context-induced nicotine-seeking is unknown, thus it will be of interest in future studies to determine this. However, because in this study our focus is on the neurobiological substrates by which contexts associated with nicotine or punishment control nicotine-seeking, we argue that the route of administration is unlikely to change the contribution of the aIC to these behaviors.

In the chemogenetics experiments, we used CLZ instead of Clozapine-N-Oxide (CNO) because CNO is converted to CLZ in-vivo (45), and made comparisons to the GFP group who also received CLZ. Various studies have estimated the conversion ratio of CNO to CLZ in rodents to be between 7.5% to 13% (46, 47), which leads to an estimate dose used here of 5-10 mg/Kg CNO, a well-established CNO dose with minimal side-effects (48, 49). While we have not controlled for non-specific effects of hM4Di expression (i.e. no vehicle-hM4Di test sessions), we find it unlikely that hM4Di expression in the absence of ligand binding will change the function of aIC given the low basal activity of this receptor (35, 50). Finally, because we used different promoters for viral-induced expression of calcium indicator (jGCamP7f: hSyn) and chemogenetic inhibition (hM4Di: mCaMKIIa), the observations reported in the photometry experiment likely reflects activity of a population of neurons that were not manipulated in the chemogenetics experiments. It will be of interest in future studies to identify potential variation in the responses of aIC neuronal subpopulations to nicotine infusions and punishment.

### Role of aIC, BLA, and BLA inputs to aIC in context-induced relapse to nicotine-seeking

In pre-clinical studies, the role of aIC in drug-seeking and relapse is well established (51–55). Bilateral electrical stimulation of the insula, at the level of mIC in this study, has been shown to decrease nicotine self-administration and both cue- and priming-induced reinstatement after extinction (56). Inactivation of both aIC and mIC can decrease nicotine self-administration, and both drug- and cue-induced reinstatement of extinguished nicotine-seeking (54, 57). Activity in aIC is also necessary for relapse to alcohol seeking, particularly relapse in the punishment context after a period of extended home cage abstinence (52). Here, we demonstrated a role for aIC in context-induced relapse to nicotine-seeking after both punishment- and extinction-imposed abstinence, further demonstrating the critical role of the aIC in the control of drug seeking.

Our results also show increased activity in BLA during context-induced relapse after punishment-imposed abstinence. The role of BLA in cue- and stress-induced reinstatement of nicotine-seeking has been demonstrated previously (58–61), but to our knowledge this is the first time that BLA has been implicated in context-induced relapse of nicotine-seeking. We also show increased activation in the aBLA→aIC pathway during context-induced relapse. Projections from BLA to aIC are critical for the maintenance of rewarding contextual stimuli (62). It has been proposed that the more posterior IC regions contribute strongly to the function of the aIC (63). However, we did not observe any increased activity in mIC→aIC or pIC→aIC neurons (either ipsi- or contra-lateral) in rats tested for context-induced relapse. Given that in this experiment, the nicotine-associated context promotes nicotine-seeking, we propose that the motivational significance of the nicotine-associated context is likely mediated through increased activity in BLA→aIC neurons and not mIC or pIC neurons.

We also found increased Fos in contralateral aIC projections during context-induced relapse. Cortico-cortical pathways are primarily thought to result in feed-forward inhibition through targeting of the PV interneurons in the contralateral hemisphere (64, 65). The contribution of contralateral cortico-cortical projections to the functions of the frontal cortex is poorly understood, and the importance of this pathway in the regulation of context-induced nicotine-seeking studied is likewise undetermined.

### Role of aIC in the regulation of nicotine taking, punishment, and seeking

Our results using fiber photometry revealed increased aIC neural activity after nose-pokes in all phases of the task. While this suggests that aIC activity may be generally important in encoding response-outcome contingencies, we observed important differences in the patterns of activity in the different phases. In self-administration, aIC activity was highest following nose-pokes that lead to nicotine infusions. During punishment, the response to nicotine infusions changed such that there was no longer a difference between nicotine reinforced nose-pokes and non-reinforced active nose-pokes. In contrast aIC activity in response to the punished nose-pokes was significantly higher than all other outcomes. Such a change in the response of aIC to (non-punished) nicotine infusion in punishment may reflect a re-evaluation of nicotine reward encoding within aIC because the punishment overcomes the motivation for nicotine and suppresses nicotine seeking. The observed reduction in nicotine seeking in punishment is an adaptive response, thus we propose that during operant behavior aIC activity may be involved in the adaptation of behavior in response to both rewarding (i.e. nicotine) and aversive outcomes (i.e. punishment). It will be of interest in future studies to determine whether maladaptive responses to punishment, such as the punishment-resistance phenotype (66), are associated with differences in the response to either reward or punishment in aIC. Interestingly, previous studies in alcohol trained rats have identified a critical role for aIC in punished alcohol seeking (67).

The insular cortex is known to support a general function of integrating interoceptive information (68), which can play an important role in addictive behaviors. Interoceptive information and external signals from environmental cues converge at the anterior insula (69, 70). In human clinical studies, insula activity is related to both positive and negative emotional reactivity. For example nicotine-associated cue exposure increases aIC activity in nicotine addicted individuals (24, 27, 71, 72), and aversive motivational states associated with short-term nicotine withdrawal are also linked to changes in resting state functional connectivity between the insula and associated brain regions (73, 74). Furthermore in non-addicted humans, insula activity is associated with punishment in a risky decision-making task (75). Positive and negative valence signals are integrated in the insula to guide motivated behavior through increased activity in the divergent outputs of insula to various brain regions (68, 70, 76–82). It will be of interest in future studies to determine whether the activity related to both positive (nicotine infusion) and negative (punished nicotine infusion) outcomes recorded at the population level in our study are selectively encoded through different output pathways of aIC.

Finally, we also found that aIC activity increased relative to baseline prior to an active nose-poke, but not inactive nose-pokes. We are unsure about the significance of this, however this pattern of activity is consistent throughout the experiment. Other studies have shown that activity in aIC is necessary for the performance of goal-directed behavior (83, 84), but it is not necessary for initial acquisition (85). We propose that the activity prior to the response differentiating between active and inactive nose-pokes may indicate that aIC activity can contribute to the encoding of expectations in operant behavior. Which may be consistent with a broader role of IC in the prediction of bodily states (70, 86)

### Similarities and differences in the neural control of relapse after punishment vs extinction

Distinct learning mechanisms are responsible for behavioral control after extinction or punishment. Both are mediated by new context-dependent associations (15, 87). However, punishment learning involves the acquisition of an association between the response and a novel outcome (shock), while extinction involves the acquisition of an association between the response and no outcome. Here we show that bilateral chemogenetic inhibition of aIC decreases context-induced relapse of both punished and extinguished nicotine-seeking. Previous studies investigating relapse of either alcohol or cocaine seeking have identified differences in the mechanisms of relapse after punishment or extinction. For example, while dopamine receptor activation in nucleus accumbens core is critical for context-induced relapse of punished alcohol seeking (18, 19), no increase in Fos was observed in nucleus accumbens core after context-induced relapse of extinguished alcohol seeking (88, 89). Inactivation of basolateral amygdala potentiates cocaine seeking after punishment, but decreases cocaine seeking after extinction (21). Meanwhile, inactivation of central amygdala had no effect on cocaine seeking after punishment, but decreased cocaine seeking after extinction (21). We propose that this finding demonstrates the importance of the aIC in context-induced relapse, regardless of the method used to impose abstinence. It has previously been demonstrated that inhibiting aIC decreases relapse of methamphetamine seeking after choice-based voluntary abstinence (55). As such the role of aIC in relapse is likely broader than just for context-induced relapse of nicotine-seeking. It will be of interest in future studies to determine whether this distinction holds true for other drugs of abuse, or indeed for other types of voluntary abstinence such as choice for social reward (90).

### Concluding remarks

We sought to investigate the neural substrates of context-induced relapse to nicotine-seeking after punishment-imposed abstinence. Our results show that activity in aIC is necessary for context-induced relapse of punished nicotine-seeking, and this is also the case for extinguished nicotine-seeking. We also show that the BLA projections to aIC are activated during context-induced relapse, and future studies are needed to determine the extent to which this activity is necessary for this relapse. Our findings further highlight the critical importance of the anterior insular cortex as a target for nicotine addiction treatments.

## Materials and Methods

### Subjects

We obtained 91 Wistar rats (29 male and 62 female), aged 10-12 weeks upon arrival, from Charles River Laboratories B.V. (Leiden, The Netherlands). In compliance with Dutch law and Institutional regulations, all animal procedures were approved by the Centrale Commissie Dierproeven (CCD) and conducted in accordance with the Experiments on Animal Act. Experiments were approved by the local animal welfare body Animal Experiments Committee of the Vrije Universiteit, Amsterdam, The Netherlands. Behavioral tests were conducted during the dark phase of the rat’s diurnal cycle (12h/12h). Food and water were available ad libitum, and rats were single-housed the rats after surgery for the remainder of the experiment.

We did not make a specific power analysis to determine sample size prior to any experiments. The group size was chosen based on our past research (7) suggesting that it will be sufficient to observe significant effects of the role of context on nicotine seeking. Each experiment is comprised of data from at least one replication cohort, and cohorts were balanced by viral group, sex, prior to the start of the experiment. We allocated the rats randomly to one of the groups within each experiment, but we were not blinded to the specific group because we were required to administer virus. We did not exclude any rats for reasons of behavioral variation (i.e. no outliers have been removed), but rats that did not have correct placement of CTb injection, or expression of jGCaMP or DREADD, within anterior insula were removed from the experiment

### Apparatus

All procedures were performed in standard Med Associates operant chambers with data collected through the MED-PC IV program (Med Associates, Georgia, VT, USA). Each chamber had one “active” and one “inactive” nose-poke hole on one wall and a grid floor connected to shock controllers. Contexts A and B were defined by houselight (on/off), cue-light color (white/red), and white noise (on/off).

The catheter for intravenous nicotine delivery was composed of a cannula connector pedestal (Plastics One, Minneapolis, MN, USA), attached to a 95 mm silicone catheter (BC-2S; 0,3 mm x 0,6 mm; UNO B.V., Zevenaar, The Netherlands) and a 6 mm piece of polyethylene tubing (0,75mm × 1,45mm; UNO B.V., Zevenaar, The Netherlands) clamping the silicone catheter to the connector pedestal. A small ball of silicone (RTV-1 Silicone Rubber / Elastosil ^®^) is attached 38 mm from the end of the silicone catheter.

For fiber photometry, excitation and emission light was relayed to and from the animal via optical fibre patch cord (0.48 NA, 400 µm flat tip; Doric Lenses). Blue excitation light (490 or 470nm LED [M490F2 or M470F2, Thorlabs]) was modulated at 211 Hz and passed through a 460-490nm filter (Doric Lenses), while isosbestic light (405nm LED [M405F1, Thorlabs]) was modulated at 531 Hz and passed through a filter cube (Doric Lenses). GCaMP7f fluorescence was passed through a 500-550nm emission filter (Doric Lenses) and onto a photoreceiver (Newport 2151). Light intensity at the tip of the fiber was measured before every training session and kept at 21uW. A real-time processor (RZ5P, Tucker Davis Technologies) controlled excitation lights, demodulated fluorescence signals and received timestamps of behavioural events. Data was saved at 1017.25Hz and analyzed with custom-made Matlab scripts.

### Drugs

Nicotine (nicotine hydrogen tartrate salt, Sigma-Aldrich, St. Louis, MO, USA) was dissolved in normal saline, filtered, and pH-adjusted to 7.4. Clozapine was dissolved first in a small amount of glacial acetic acid (volume used was 0.1% of the final CLZ volume) and progressively diluted in saline until a final concentration of 0.3 mg/ml (pH was adjusted to 7.0 – 7.2).

### Viral vectors

We purchased premade viral vectors from the University of Zurich viral vector core: AAV-5/2-mCaMKIIa-HA_hM4D(Gi)-IRES-mCitrine-WPRE-hGHp(A) (**hM4Di**), AAV-5/2-mCaMKIIa-EGFP-WPRE-hGHp(A) (**GFP**), AAV-9/2-hSyn1-chI-jGCaMP7f-WPRE-SV40p(A) (**jGCaMP7f**). The titer injected was: hM4Di, 2.4×10^12 gc/ml; GFP, 2.5×10^12 gc/ml; jGCaMP7f, 4.4×10^12 gc/ml.

### Surgery

Thirty minutes prior to surgery, we injected rats with the analgesic Rymadil^®^ (5 mg/kg; Merial, Velserbroek, The Netherlands) and the antibiotic Baytril^®^ (8.33 mg/kg; Bayer, Mijdrecht). Surgery was performed under isoflurane gas anesthesia (PCH; Haarlem). The silicone catheter was tunneled from the scalp to the neck and was inserted into the jugular vein, where it was secured using sterile thread. We sealed the silicone catheter using a taurolidine-citrate solution (TCS; Access Technologies, Skokie, IL) and a polyethylene cap. After the catheter was implanted, we placed the rat in a stereotactic frame (David Kopf Instruments, Tujunga, CA) and injected Xylocaïne 2% with adrenaline (10 mg/kg; Astra Zeneca, Zoetermeer, The Netherlands) into the incision site prior to the incision. A craniotomy above aIC was performed, followed by CTb or AAV injections (see below for details). After filling the skull hole with bone wax, cannula tubing connected to a Plastics One Connector-Pedestal and optic fiber implant (when applicable) was secured to the skull using dental cement (IV Tetric EvoFlow 2g A1, Henry Schein, Almere) and jewelers screws. Rymadil (5 mg/kg; s.c.) was administered for 2 days after the surgery. Rats were given one week of recovery following surgery.

#### CTb injections

40 nl of 1% CTb (List Biological Laboratories) was injected unilaterally (left or right) into aIC (AP: +2.8, ML: +4.0, DV: −5.9 mm from Bregma) over 2 min using 1.0 μl 32 gauge “Neuros” syringe (Hamilton) attached to a UltraMicroPump (UMP3) with SYS-Micro4 Controller (World Precision Instruments). The needle was left in place for an additional 2 min after injections.

#### AAV injections for fiber photometry

0.5 μl of AAV solution was injected unilaterally (left or right) into aIC (AP: +2.8, ML: +4.0, DV: −6.0 mm from Bregma) over 5 min. The needle was left in place for an additional 5 min. A 400μm optic fiber (Doric Lenses) was then implanted above aIC (AP: +2.8, ML: +4.0, DV: −5.6 mm from Bregma).

#### AAV injections for chemogenetics

1.0 μl of AAV solution was injected bilaterally into aIC (AP: +2.8, ML: +4.0, DV: −5.9 mm from Bregma) over 5 min using 10 μl Nanofil syringes (World Precision Instruments), with 33 gauge needles, attached to a UltraMicroPump (UMP3) with SYS-Micro4 Controller (World Precision Instruments). The needle was left in place for an additional 5 min.

### Behavioral procedure

#### Phase 1: Nicotine self-administration (SA: Context A)

On the day prior to self-administration, before and after each self-administration session and during weekends, rat’s catheters were flushed with approx. 0.1 ml mixture of heparin (0.25 mg/ml; Serva, Heidelberg, Germany) and gentamicin sulfate (0.08 mg/ml; Serva, Heidelberg, Germany). Rats were trained to self-administer nicotine in 2-hour sessions, five days a week. Entry into the active nose-poke resulted in intravenous nicotine delivery infused over approximately 2 seconds (infusion time adjusted for weight) at 30 μg/kg/infusion. Nicotine infusion was paired with a 20-second time-out period with the cue-light on. During time-out, responses were recorded but had no consequence. Inactive nose-pokes had no effect in either context. Rats were first trained on a fixed-ratio (FR) 1 schedule, which was then increased to FR-2, followed by FR-3. We tested catheter patency using intravenous anesthetic 0.05 cc pentothal (thiopenthal sodium, 50 mg/ml).

#### Phase 2A: Punishment of nicotine self-administration (PUN, Context B)

Nicotine self-administration was maintained on the FR-3 schedule, and 50% of the reinforced active nose-pokes (pseudo-randomly determined by the Med-PC program) resulted in footshock (0.30 mA for 0.5 sec) and nicotine infusion.

#### Phase 2B: Extinction of nicotine-seeking (EXT, Context B)

Entry into the active nose-poke resulted in simultaneous activation of the cue light and delivery of the same volume of saline through the jugular catheter (FR-3 schedule).

#### Phase 3: Relapse tests in Context A (SA) & Context B (PUN or EXT)

Following abstinence (PUN or EXT), rats were tested in context A and context B. The response contingent CS was presented on an FR-3 schedule without punishment, saline or nicotine delivery. For the CTb+Fos experiment (Exp. 1), rats were tested for one 60 minute session and perfused 90 minutes after the beginning of the test. For the chemogenetics experiments (Exp. 3 and 4) rats were tested in both contexts A and B (counterbalanced order). 30 min prior to the relapse test sessions, we injected both GFP and hM4Di expressing rats with clozapine at a dose of 0.3 mg/kg injection (i.p.).

### Immunohistochemistry

We deeply anesthetized rats with isoflurane and Euthasol^®^ injection (i.p.) and transcardially perfused them with ~100 ml of normal saline followed by ~400 ml of 4% paraformaldehyde in 0.1M sodium phosphate (pH 7.4). The brains were removed and post-fixed for 2 h, and then 30% sucrose in 0.1M PBS for 48 h at 4°C. Brains were then frozen on dry ice, and coronal sections were cut (40 μm) using a Leica Microsystems cryostat and stored in 0.1M PBS containing 1% sodium azide at 4°C.

Immunohistochemical procedures are based on our previously published work (17, 19, 89). We selected a 1-in-4 series and first rinsed free-floating sections (3 x 10 minutes) before incubation in PBS containing 0.5% Triton-X and 10% Normal Donkey Serum (NDS) and incubated for at least 48 h at 4°C in primary antibody. Sections were then repeatedly washed with PBS and incubated for 2-4 h in PBS + 0.5% Triton-X with 2% NDS and secondary antibody. After another series of washes in PBS, slices were stained with DAPI (0.1 ug/ml) for 10 min prior to mounting onto gelatin-coated glass slides, air-drying and cover-slipping with Mowiol and DABCO.

#### CTb+Fos protein labeling

Primary antibodies were rabbit anti-c-Fos (1:2000; Cell Signaling, CST5348S) and goat anti-CTb (1:5000; List Biological Laboratories, 703). Secondary antibodies were donkey anti-rabbit Alexa Fluor 594 (1:500; Molecular Probes, A21207) and donkey anti-goat Alexa Fluor 488 (1:500: Molecular Probes, A11055).

#### Photometry experiment

Primary antibodies were mouse anti-NeuN primary antibody (1:1000; Chemicon, MAB377) and rabbit anti-GFP primary antibody (1:2000; Chemicon, AB3080). Secondary antibodies were donkey anti-mouse DyLight 649 (1:500; Jackson ImmunoResearch, 715-495-150) and donkey anti-rabbit Alexa Fluor 594 (1:500: Mol. Probes, A21207).

#### Chemogenetic inhibition experiments

Primary antibody was rabbit anti-GFP primary antibody (1:2000; Chemicon, AB 3080) and secondary antibody was donkey anti-rabbit Alexa Fluor 594 (1:2000; Molecular Probes, A21207). In rats in the GFP group, slices were only stained with DAPI.

### Image acquisition and neuronal quantification

Slides were all imaged on a VectraPolaris slide scanner (VUmc imaging core) at 10x magnification. For the CTb+Fos experiment (Exp. 1), images from Bregma +4.0 to Bregma −3.3 were scanned and imported into QuPath for analysis (91). For photometry and chemogenetic experiments, images containing aIC, from Bregma + 4.2 mm to +2.5 mm were identified and the boundary of expression for each rat was plotted onto the respective Paxinos and Watson atlas (92). Rats in Experiment 1 that had a CTb injection not within aIC were excluded from analysis. Rats in Experiments 4 and 5 that had either unilateral expression or misplaced expression were excluded from the analysis.

#### Fos, CTb, and CTb+Fos quantification

Regions of interest were manually labeled across sections using DAPI for identification of anatomical landmarks and boundaries. For the IC, we labeled 18 sections per hemisphere per rat spaced approximately 400 microns apart. For analysis, we separated IC into three regions anterior (aIC), middle (mIC), and posterior (pIC), and each value was the result of an average of each count from six adjacent sections: aIC (approx. Bregma +3.72 to +1.44), mIC (approx. Bregma +1.08 to −0.72), pIC (approx. Bregma −1.08 to −2.92). For BLA, we labeled 6 sections spaced approximately 200 microns apart, and we separated it into anterior BLA (aBLA) and posterior BLA (pBLA), which is the average value of three adjacent sections: aBLA (approx. Bregma −1.92 to −2.40), pBLA (approx. Bregma −2.64 to −3.12). Some rats had missing sections due to mistakes during the process, and these sections were left blank for the statistical analyses.

To identify Fos- and CTb-positive cells, we used the ‘Cell detection’ feature in QuPath, with an identical threshold applied across all sections. CTb was not counted for the first six sections in the ipsilateral aIC, where the CTb injection was located, because the cell detection feature could not reliably discriminate between CTb positive cell and the CTb injection. The total number of positive cells per region was divided by the area in mm2. To identify CTb+Fos cells, each region of interest was exported to ImageJ. The overlays representing the cells (CTb or Fos) were then filled, converted to a binary layer, and then multiplied using the ImageJ function ‘Image calculator’. The nuclei that remained as a result of this function were counted as double-labeled CTb+Fos neurons. CTb+Fos double labelling is reported as a percentage of total CTb neurons for that given region of interest.

### Ex-vivo slice physiology

Coronal slices were prepared for electrophysiological recordings. Rats were anesthetized (5% isoflurane, i.p. injection of 0.1 ml/g pentobarbital) and perfused with ice-cold N-Methyl-D-glucamin (NMDG) solution containing (in mM): NMDG 93, KCl 2.5, NaH2PO4 1.2, NaHCO3 30, HEPES 20, Glucose 25, sodium ascorbate 5, sodium pyruvate 3, MgSO472H2O 10, CaCl2*2H2O 0.5, at pH 7.3 adjusted with 10 M HCl. The brains were removed and incubated in ice-cold NMDG solution. 300 μm thick brain slices were cut in ice-cold NMDG solution and subsequently incubated for 15-30 min at 34 °C.

Before the start of experiments, slices were allowed to recover for at least 1 hour at room temperature in carbogenated (95% O2/5% CO2) ACSF solution containing (in mM): NaCl 125, KCl 3, NaH2PO4 1.2, NaHCO3 25, Glucose 10, CaCl2 2, MgSO4 1. For voltage- and current-clamp experiments borosilicate glass patch-pipettes (3–5 MΩ) were used with a K-gluconate-based internal solution containing (in mM): K-gluconate 135, NaCl 4, MgATP 2, Phosphocreatine 10, GTP (sodium salt) 0.3, EGTA 0.2, HEPES 10 at a pH of 7.4. Data was sampled using a Multiclamp 700B amplifier (Axon Instruments) and pClamp software (Molecular Devices). All recordings were made between 31.1°C and 33.6°C.

### Experimental design

#### Exp. 1: CTb + Fos after context-induced relapse of punished nicotine-seeking (n=18 female. Cohort 1, n=11; cohort 2, n=7)

Fig. 1A shows the experimental outline. We first trained rats to self-administer nicotine in one context (context A) in 2 h sessions per day for 15 days (4 sessions FR-1, 6 sessions FR-2, 5 sessions FR-3). Next, the rats underwent punishment in the alternate context (context B) 2 h per day for 7 days. During these sessions, active nose-pokes (FR-3 schedule) resulted in the presentation of the cue-light (20 sec), nicotine infusion, and 50% probability of 0.3 mA footshock punishment. Finally, the rats were tested under extinction conditions. One group of rats (n=7) was tested in the nicotine self-administration context (context A), one group (n=8) was tested in the punishment context (context B), and a third group (n=3) was taken from the home-cage without test. We perfused rats 90 min after the start of the 60 minute test.

#### Exp.2: Calcium imaging of aIC activity during nicotine self-administration, punishment, and context-induced relapse (n = 7 female. Cohort 1, n=3; cohort 2, n=4)

Fig. 2A shows the experimental outline. Photometry sessions were 1 hour in duration. Tubing delivering nicotine was run down along the patch-cord and to the connection in the implant. If the tubing became tangled, the rat was manually rotated in the opposite direction or, if it was within 10 minutes of the end of the session, the session was ended early. Rats were trained to self-administer nicotine in context A (7 days FR-1, 3 days FR-2, 8 days FR-3). We next punished nicotine self-administration in context B for 6 days (first cohort; n=3) or 10 days (second cohort; n=3), one rat only received 4 days of punishment. In this phase 50% of nicotine infusions were paired with electric shock. After punishment-imposed abstinence, nicotine-seeking was tested in context A in 2 x 1 hour sessions over 2 consecutive days.

#### Exp. 3: Chemogenetic validation (n = 3 female, n = 3 male)

Six rats were unilaterally injected with 1.0 μl of AAV encoding the inhibitory DREADD hM4Di into aIC (AP: +2.8, ML: +4.0, DV: −5.9 mm from Bregma). Rats were sacrificed 4-5 weeks later for ex-vivo physiology.

#### Exp. 4: Effect of chemogenetic inhibition of aIC on context-induced relapse of punished nicotine-seeking (n=28 (13M/15F). Cohort 1, n=17 (8M/9F); cohort 2, n=11 (5M/6F))

Fig. 4A shows the experimental outline. We first trained rats to self-administer nicotine in context A (4 days FR-1, 4 days FR-2, 5 days FR-3). We next punished nicotine self-administration in context B for 6 days. We then tested rats in both contexts (A and B), over 2 consecutive days, and the order was counterbalanced. 30 min prior to the test session, we injected both GFP and hM4Di expressing rats with clozapine (0.3 mg/kg injection (i.p.). We excluded 3 female rats from the hM4Di group because of a lack of bilateral hM4Di expression in aIC.

#### Exp. 5: Effect of chemogenetic inhibition of aIC on context-induced relapse of extinguished nicotine-seeking (n=31 (13M/18F). Cohort 1, n=22 (11M/11F); cohort 2, n=9 (2M/7F))

Fig. 5A shows the experimental outline. We first trained rats to self-administer nicotine in context A on FR-1 (4 days), then FR-2 (4 days), then FR-3 (5 days). Next, we extinguished nicotine-seeking by saline infusion in context B (EXT) for 8 days. We then tested rats in both contexts (A and B), over 2 consecutive days, and the order was counterbalanced. 30 min prior to the test session, we injected both GFP and hM4Di expressing rats with clozapine (0.3 mg/kg injection (i.p.).

### Statistics

All behavioral data was analyzed using IBM SPSS V21. Phases were analyzed separately. Dependent variables were the total number of active and inactive nose-pokes across phases, and nicotine infusions for nicotine self-administration and punishment phases. For the CTb+Fos test (Exp. 1) we used a repeated measures analysis of variance (ANOVA) with Nose-Poke (Active, Inactive) as a within-subjects factor and Test Context (context A, context B) as the between-subjects factor. To analyze CTb+Fos expression, we used one-way ANOVA to test for an effect of Test Context (Home-cage, Punishment, Nicotine) on Fos, CTb, and % CTb+Fos/CTb. Follow-up tests (Tukey) were conducted on regions that had a significant main effect of Test Context. For the chemogenetic experiments we used repeated measures ANOVA with Test Context (context A, context B) and Nose-Poke (Active, Inactive) as within-subjects factors, and Virus (GFP, hM4Di) and Sex (Female, Male) as between-subjects factors.

#### Photometry

Recorded signals were first downsampled by a factor of 64, giving a final sampling rate of 15.89 Hz. The 405nm isosbestic signal was fit to the 490nm calcium-dependent signal using a first order polynomial regression. A normalized, motion-artefact-corrected ΔF/F was then calculated as follows: ΔF/F = (490nm signal − fitted 405nm signal)/fitted 405nm signal. The resulting ΔF/F was then detrended via a 90s moving average, and low-pass filtered at 3Hz. ΔF/F from 5s before nosepoke (baseline) to 10s after nosepoke were collated. These traces were then baseline-corrected and converted into z-scores by subtracting the mean baseline activity during first 4 seconds of the baseline and dividing by the standard deviation of those 4 seconds. To avoid duplicate traces due to overlapping epochs, we excluded from the analyses any nosepokes that occurred within 20s after a rewarded nosepoke (time-out), and un-rewarded active/inactive nosepokes that occurred 5s after another active/inactive nosepoke.

Nosepoke traces were grouped by response type: active rewarded, active punished, active non-rewarded, and inactive. Two analysis approaches were used, bootstrapping and permutation tests, the rationale for each is described in detail in (38).

Bootstrapping was used to determine whether calcium activity per response type was significantly different from baseline (ΔF/F = 0). A distribution of bootstrapped means were obtained by randomly sampling from traces with replacement (*n* traces for that response type; 5000 iterations). A 95% confidence interval was obtained from the 2.5^th^ and 97.5^th^ percentiles of the bootstrap distribution, which was then expanded by a factor of sqrt(n/(n – 1)) to account for narrowness bias (38).

Permutation tests were used to assess significant differences in calcium activity between response types. Observed differences between response-types were compared against a distribution of 1000 random permutations (difference between randomly regrouped traces) to obtain a p-value per time point. Alpha of 0.05 was Bonferroni-corrected based on the number of comparison conditions, resulting in alpha of 0.01 for comparisons between self-administration and test sessions (3 conditions) and alpha of 0.008 for punishment sessions (4 conditions). For both bootstrap and permutation tests, only periods that were continuously significant for at least 0.25s were identified as significant (38).

## Funding and Disclosure

The work was supported by a Fulbright Fellowship to RH, and Austrian Science Fund (FWF) grant SPIN supportW1206-12 to HG and GZ (graduate program Signal Processing In Neurons, www.neurospin.at). The authors declare no conflict of interest.

## Acknowledgments

The authors gratefully acknowledge the Histology Imaging Unit for their support & assistance in whole-slide imaging. The authors would like to thank Francesco Ferraguti for insightful comments in the preparation of this manuscript.

## Author contributions

Conducted experiments: HG, IAL, RH, DS, YvM, TH, NJM. Analyzed data: HG, IA, RH, PJRDB, TH, NJM. First draft of manuscript: HG, IAL, NJM. Edited subsequent drafts and finalized manuscript: HG, IAL, NJM, PJRDB, GZ, HM, TdV.

## Notes

### Competing Interest Statement

The authors have declared no competing interest.

